# Shorter self-reported sleep duration in men is associated with worse virtual spatial navigation performance

**DOI:** 10.1101/2023.09.09.556942

**Authors:** Emre Yavuz, Alpar S Lazar, Christoffer J. Gahnstrom, Sarah Goodroe, Antoine Coutrot, Michael Hornberger, Hugo J. Spiers

## Abstract

Sleep has been shown to impact navigation ability. However, it remains unclear how different sleep-related variables may be independently associated with spatial navigation performance, and as to whether gender may play a role in these associations. We used a mobile video game app, Sea Hero Quest (SHQ), to measure wayfinding ability in US-based participants. Wayfinding performance on SHQ has been shown to correlate with real-world wayfinding. Participants were asked to report their sleep duration, quality, daytime sleepiness and nap frequency and duration on a typical night (n = 737, 409 men, 328 women, mean age = 27.1 years, range = 18-59 years). A multivariate linear regression was used to identify which self-reported sleep variables were independently associated with wayfinding performance. For men, longer self-reported sleep durations were associated with better wayfinding performance. For women, no such association was found. While other self-reported sleep variables showed trends of association with wayfinding performance, none of these were significantly associated with wayfinding performance in our regression model. These findings from younger U.S.-based participants suggest that a longer self-reported sleep duration may be an important contributor to successful navigation ability in men.

## Introduction

Being able to maintain a sense of direction and location in order to find our way in different environments is a fundamental cognitive function that relies on multiple cognitive domains^1, 2^. Understanding these individual differences in navigation ability is crucial given that deficits in navigation may constitute the earliest signs of Alzheimer’s Disease^3^. There are also negative effects of disorientation, such as getting lost in everyday environments, leading to distress for patients and family members and in extreme cases death from exposure^4^. Understanding these individual differences in navigation will also advance the field of spatial cognition at large^1, 2^. Being able to create a valid test of navigation that accounts for the wide variation in navigation performance is challenging, given the large sample sizes needed and the high levels of environmental manipulation and experimental control required in standard research settings^1^.

With the recent evolution of widespread touch-screen technology on both tablet and mobile devices and virtual reality (VR), our team developed a series of navigation tests in the form of a mobile video game app Sea Hero Quest (SHQ)^2^. SHQ has since been used to test the navigation ability of 4 million people globally, has good test-retest reliability and is predictive of real-world navigation performance^4–6^.

Sleep quality has previously been associated with spatial navigation performance, where both subjective and objective measures of sleep quality have been linearly associated with poorer virtual spatial navigation performance^7–9^, including sleep duration^7^, fragmented sleep^7, 8^ and insomnia-like symptoms^8^. Experimental sleep deprivation studies have also shown that sleep deprivation results in poorer spatial navigation performance^10–13^, although other studies report no significant changes in spatial navigation performance following sleep deprivation^14, 15^. A recent study from our team also reported a quadratic (U-shaped) association between self-reported sleep duration and virtual spatial navigation performance on SHQ, where mid-range sleep duration was associated with optimal performance^16^. This latter study corroborates broader findings of a U-shaped association between self-reported sleep duration and other cognitive domains aside spatial navigation^17, 18^, even when controlling for other sleep-related variables such as daytime sleepiness, sleep apnea and insomnia symptoms^19, 20^. However, it remains unclear as to how different sleep-related variables may independently be associated with spatial navigation performance when accounting for other sleep-related variables.

Gender differences have also been found in various aspects of sleep, with poorer self-reported sleep quality in women^19, 21–23^ and shorter self-reported and objective sleep duration in men^16, 24^. The association between self-reported duration and cognitive function have also been shown to differ by gender, where shorter and longer sleep durations are associated with worse cognitive performance only in men^25^. Gender differences have also been found in both virtual and real-world navigation ability in favour of males^2, 5, 26^. Despite these findings, no studies have examined how the association between sleep and human spatial navigation performance may differ by gender.

With both poorer sleep quality and poorer spatial navigation performance being associated with a higher risk of developing various neurodegenerative disorders^3, 27–28^, a deeper understanding of the association between sleep and spatial navigation performance at the population level will provide greater insight into the factors that may contribute to cognitive decline.

Bringing together these findings, we hypothesised that self-reported sleep duration, time spent awake during the night and sleep quality would show a linear association with human spatial navigation performance, where those with shorter self-reported sleep duration, greater self-reported time spent awake during the night and a poorer self-reported sleep quality would show worse spatial navigation performance. Based on a previously shown quadratic (inverted U-shaped) association between sleep duration and spatial navigation performance^16^, we also hypothesised that sleep duration would show a quadratic association with human spatial navigation performance. We therefore specified sleep duration as both a linear and quadratic term. Given the previously found gender differences in both sleep-related variables and human spatial navigation performance, we also hypothesised that gender would show a significant interaction with self-reported sleep duration (both when expressed as a linear and quadratic term), self-reported time spent awake during the night and self-reported sleep quality, but did not specify a direction for this interaction. In addition to our hypotheses, we explored whether other self-reported sleep-related variables aside sleep duration would be independently associated with navigation ability and as to whether these associations would differ by gender.

## Methods

### Participants

903 participants living in the US aged 18 and above (365 men, 459 women, mean age = 27.1 years, SD = 8.0 years, range = 18-66 years, mean number of years spent in formal education = 16.1 years, mean BMI = 26.4, 285 currently living cities, 556 currently living outside cities) were recruited using the Prolific database^29^ and reimbursed for their time. Ethical approval was obtained from the University College London Review Board conforming to Helsinki’s Declaration. All participants provided informed written consent. We removed 17 participants who selected ‘other’ for gender, as we were primarily interested in the differences in the associations between the sleep variables and wayfinding distance in men and women. We then excluded outliers in 2 stages. We first removed participants whose time spent in bed was less than 4 hours and greater than 12 hours, as we believed these participants’ responses were not genuine ot considered them to be either extremely short or extremely long sleepers respectively, based on a criteria used by a previous study^30^. This resulted in the removal of 62 participants, and a remaining sample of 824 participants. We then further identified outliers in this remaining sample of 824 participants using Mahalanobis’ Distance, a method shown to have high sensitivity and specificity and a minimal change in bias in simulated and real datasets when removing outliers based on questionnaire data compared to other methods^31, 32^ (Supplementary Materials). 97 participants were removed as outliers. This resulted in a final sample of 724 participants (328 men, 409 women, mean age = 26.4 years, SD = 7.2 years, range = 18-59 years, mean number of years spent in formal education = 16.1, mean Body Mass Index (BMI) = 26.0, 248 currently living cities, 476 currently living outside cities). Demographic information for the final sample is summarised in Table 1. Data analysis was completed using R studio (version 1.4.1564) and Python (version 3.9.12). T-tests were used to determine whether sleep-related variables differed in men and women for the continuous sleep variables. A chi-squared analysis was used to determine whether the frequency of alarm use differed in men and women given that this was a categorical variable.

**Table 1.** Summary of basic descriptive statistics for the variables included in the model (weighted wayfinding distance, age, highest level of education achieved, weekly hours of video game use on all devices, weekly hours of phone use, BMI, daily hours of sunlight, daily cups of coffee, weekly units of alcohol and frequency of daily strenuous physical activity) in men and women separately. Mean and standard deviation (SD) values are shown. Men were significantly older than women, spent significantly greater weekly hours video gaming and had significantly fewer weekly hours of phone use.

|  | Female | Male |
| --- | --- | --- |
| n | 410 | 314 |
| Weighted wayfinding distance | 12.21 (4.03) | 11.06 (4.88) |
| Age | 25.27 (6.83) | 27.98 (7.38) |
| Highest level of education achieved (n, %) |  |  |
| Some formal education | 5 (1.2) | 2 (0.6) |
| High school | 109 (26.6) | 111 (35.4) |
| 2-year college/uni | 57 (13.9) | 43 (13.7) |
| 4-year college/uni | 178 (43.4) | 117 (37.3) |
| Masters | 55 (13.4) | 30 (9.6) |
| PhD/MD | 6 (1.5) | 11 (3.5) |
| Hours of phone use per week | 35.60 (23.08) | 28.59 (21.54) |
| Weekly hours of video gaming on all devices | 4.67 (5.23) | 9.67 (6.19) |
| BMI | 26.44 (7.24) | 25.34 (5.67) |
| Daily hours of sunlight | 1.24 (1.11) | 1.38 (1.24) |
| Daily cups of coffee | 1.12 (1.09) | 1.44 (1.48) |
| Weekly units of alcohol | 3.09 (5.14) | 3.04 (5.16) |
| Frequency of daily strenuous physical activity |  |  |
| (%) |  |  |
| Never | 9.8 | 10.5 |
| Rarely | 37.8 | 38.9 |
| Sometimes | 37.8 | 37.9 |
| Often | 12.7 | 9.9 |
| Always | 2.0 | 2.9 |
| Smoking frequency (%) |  |  |
| Never smoked | 85.1 | 72.0 |
| Former smoker | 11.5 | 16.2 |
| Current smoker | 3.4 | 11.8 |

### Statistical Power Analysis

A power analysis was conducted using G* power^33^. Given that a previous study looking at the association between sleep quality and spatial navigation ability found a medium effect size for this association^9^, it was determined that 737 participants were sufficient to acheive a medium effect size (Cohen’s f2 = 0.15) at an alpha threshold of 0.05 with 95% power^34^.

## Experimental Procedure

### Self-report questionnaires

To characterize self-reported sleep patterns, we used a series of self-report questions about bedtime, wake-up time, difficulty waking up, sleep quality, number of awakenings during the night, time spent awake during the night, sleep duration, sleepiness on waking (sleep inertia), sleepiness on going to bed, nap duration, frequency of naps, and frequency of alarm use (see Supplementary Materials). All of these variables related to a typical night of sleep. As sleep quality was originally coded in the opposite direction to the other variables (1 = maximum, 10 = minimum), we reverse coded this variable to be consistent with the other variables with the coding strategy used for the other variables we collected (1 = minimum, 10 = maximum). We also calculated two additional sleep-based measures - the sleepiness resolution index (SRI) and the time in bed (hours). The SRI was a measure developed by our research team to determine the extent to which one’s level of sleepiness changes after a typical night’s sleep. The SRI was calculated the self-reported sleepiness upon bedtime over that upon self-reported wake-up time. Thus, an SRI value of greater than 1 would indicate that one was more sleepy when going to bed than when waking up. The time in bed (hours) was calculated as the time difference between self-reported bedtime and self-reported wakeup time. We were unable to derive a measure of sleep fragmentation as we did not collect data on the length of each awake period during the night^7^.

To characterize the individual video game use, participants were asked to indicate the number of hours per week spent playing video games on all kind of devices and the number of hours of phone use per week.

Participants were also asked to report their age, gender, highest education level, height (in feet and inches), smoking frequency (currently, previous or never smoked), weekly units of alcohol intake, daily cups of caffeine, frequency of daily strenuous physical activity. BMI was calculated as weight (kg) divided by height squared (m^2).

Please see the Supplementary Materials for the full set of questionnaires.

### Sea Hero Quest Task

Sea Hero Quest (SHQ) is a VR-based video game for mobile and tablet devices which requires participants to navigate through a three-dimensional rendered world in a boat to search for sea creatures in order to photograph them, with the environment consisting of ocean, rivers and lakes^2, 5^. Although navigational abilities on SHQ have been assessed using Wayfinding, Path Integration and Radial Arm Maze measures, we focused on wayfinding in this study (Figure 1). We asked participants to play 5 levels - levels 1, 11, 32, 42 and 68 - where level 1 was a tutorial level designed to assess one’s ability to control the boat, whilst the latter 4 levels were wayfinding levels. We selected these specific levels as they showed the greatest effect sizes when investigating the effect of one’s home environment on wayfinding performance in a previous study^16^. The wayfinding levels increased in difficulty from level 11 to 68, with the difficulty of a given level based on the number of goals and how far apart they were from each other. Participants were required to play all 5 levels, where playing a given level would unlock the next level.

**Figure 1.**
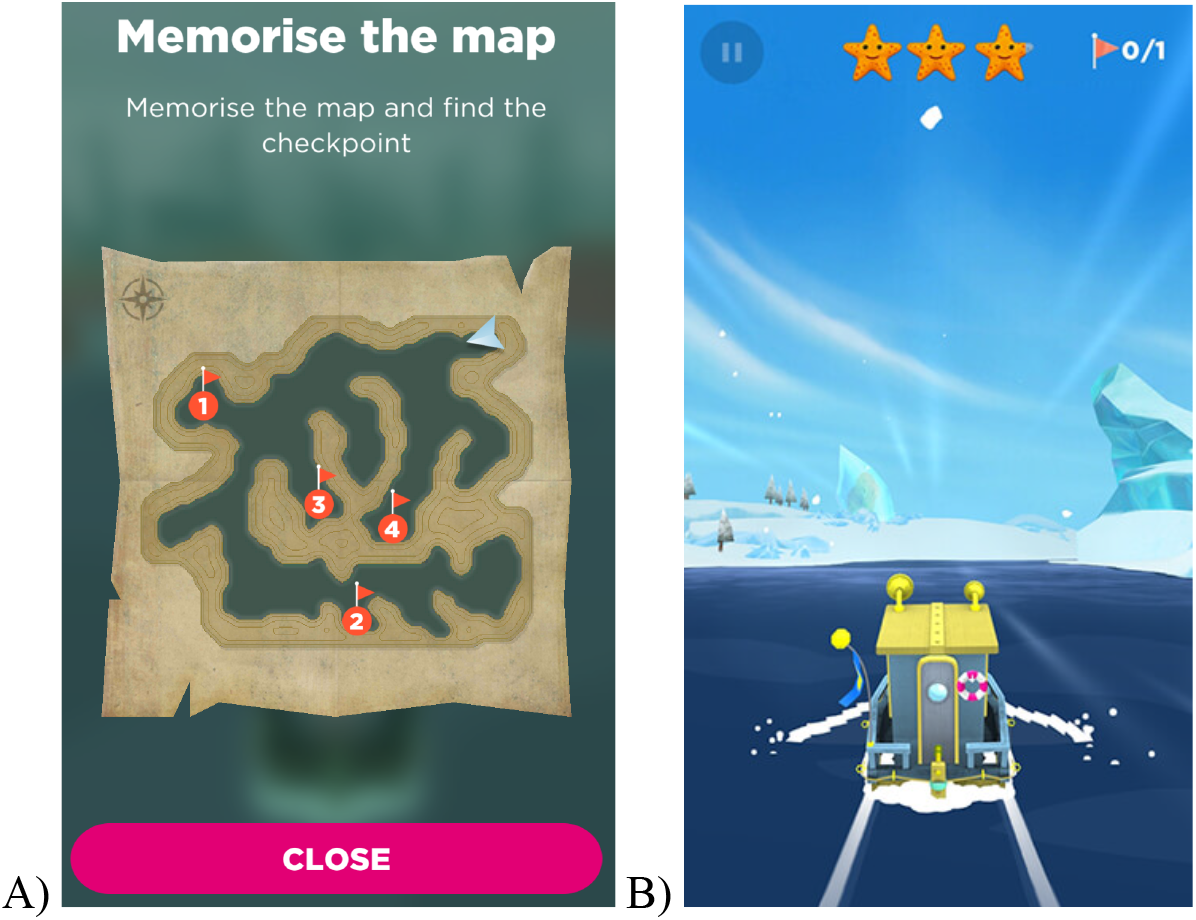
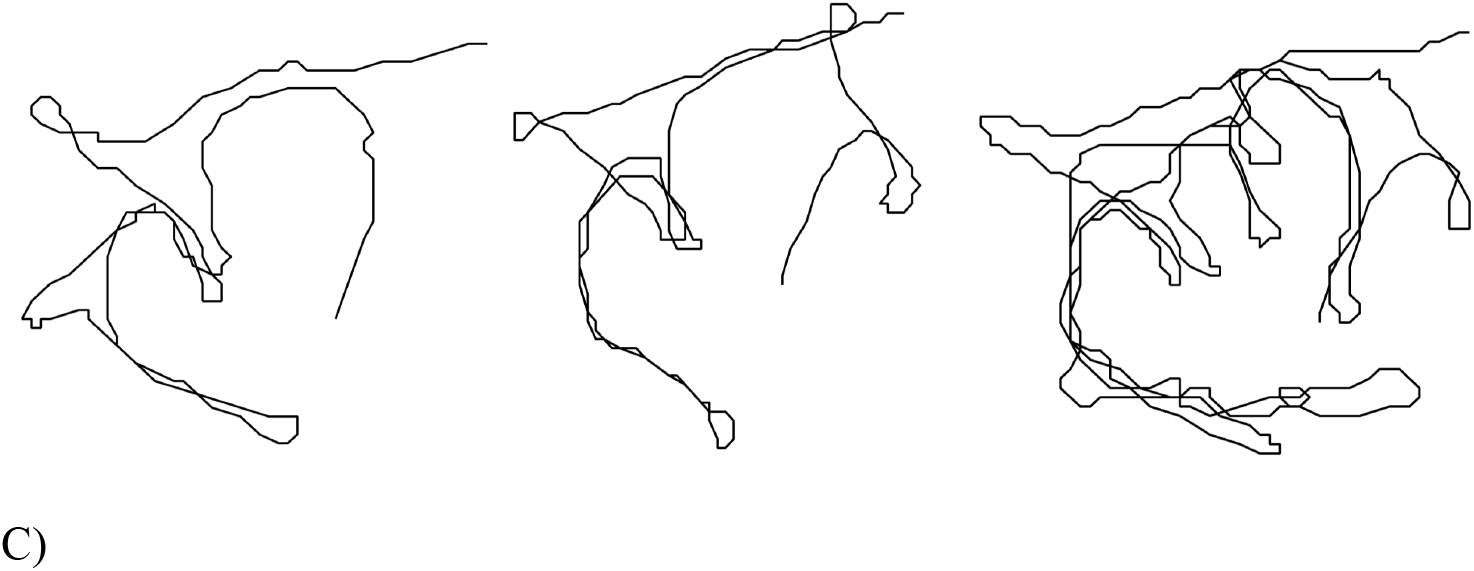
Outline of the wayfinding task. (A) At the start of each game level participants were presented with a map indicating the goals which they had to navigate to and the order of visitation, where ‘1’ indicates goal number 1 etc. (the map from level 32 is shown as an example). Level 1 (not shown) provided one goal and a simple river with no choices to reach the goal as training. (B) Participants selected to close the map by pressing ‘close’, at which point the participant had to start navigating to the goals using their memory of the map. First-person view of the environment is shown, where the participant tapped left or right of the boat to steer it. (C) Examples of individual player trajectories (level 32) from the start location to the final goal. Trajectories are ordered by performance, with the top left providing the best performance (shortest trajectory length), through to bottom right who has the worst performance (longest trajectory length). Adapted from a previous study^35^.

For each participant, we quantified the wayfinding distance, defined as the euclidean distance travelled between each sampled location in pixels, for levels 11, 32, 42 and 68 separately. We then divided the wayfinding distance in each level by the wayfinding distance in level 1 to control for the effects of gaming experience on navigation performance. To control for the difference in wayfinding distance between levels, we then z-scored the distances within each level and averaged these across the levels. This resulted in each participant having a z-score which represented their wayfinding distance across the 4 levels. This z-score was referred to as the weighted wayfinding distance. A shorter weighted wayfinding distance indicated better navigation performance (i.e., a more efficient route to the goal).

### Data Analysis

To investigate the relationship between sleep and spatial navigation, we ran partial spearman’s correlations between the continuous sleep-related variables we collected (time spent awake during the night, nap duration, hours of sleep, number of awakenings during the night, sleepiness resolution index, time in bed, difficulty waking up, sleep inertia, sleepiness on waking and sleep quality) and weighted wayfinding distance in SHQ. For the binary sleep-related variables (take naps yes/no and use alarms yes/no) and weighted wayfinding distance, we used point-biserial correlations. When running these correlations, we controlled for age, gender, highest level of education achieved, daily hours of sunlight, daily cups of caffeine, daily hours of strenuous physical activity, weekly units of alcohol and Body Mass Index (BMI). We included these covariates to control for factors associated with sleep or wayfinding distance aside the sleep-related variables^35–40^. We also included weekly hours of phone use as a covariate given that SHQ is a mobile phone-based video game.

We then entered the five self-reported sleep variables with the strongest effect sizes (i.e., the strongest association with wayfinding distance as indicated by the spearman’s correlation coefficient) into a multivariate linear regression model. This was to determine the independent associations between each of the sleep variables and navigation performance when controlling for other variables. Age, gender, highest level of education, weekly hours of video gaming, weekly hours of phone use, daily hours of sunlight, daily cups of caffeine, daily hours of strenuous physical activity and Body Mass Index (BMI) were also included as covariates in this multivariate model for the reasons mentioned above. A sleep variable x gender interaction term for each sleep variable was included in the model as we were interested in the role played by gender in the association between the sleep-related variables and navigation. Given that sleep duration has been shown to have both a U-shaped association with spatial navigation ability on SHQ^16^ and a linear association with spatial navigation ability on other tasks^7^, we specified two models. In the first model, sleep duration was included as a linear term (Table 3). In the second model, sleep duration was included as a quadratic term (Table 4).

As a check for multicollinearity, we calculated the variance inflation factor (*VIF*) for each predictor variable in each model (Supplementary Tables S1 and S2). A *VIF* value of <5 indicates that multicollinearity is not a concern^41, 42^. Post-hoc t-tests bonferroni-corrected for multiple comparisons to control for type 1 errors were carried out where main effects and interactions were significant. When conducting post-hoc t-tests for sleep duration, we binned sleep duration into <=7 hours and >7 hours of sleep to match the approach taken by other studies^17, 22, 25, 43^. Given that only 5 participants slept for more than 9 hours on a typical night, we grouped those who slept for 9 hours or 10 hours on a typical night into one group (>= 9 hours on a typical night) when visualising the data.

## Results

### Self-reported sleep-related characteristics

Please see Table 2 for self-reported sleep-related characteristics in men and women.

**Table 2.**
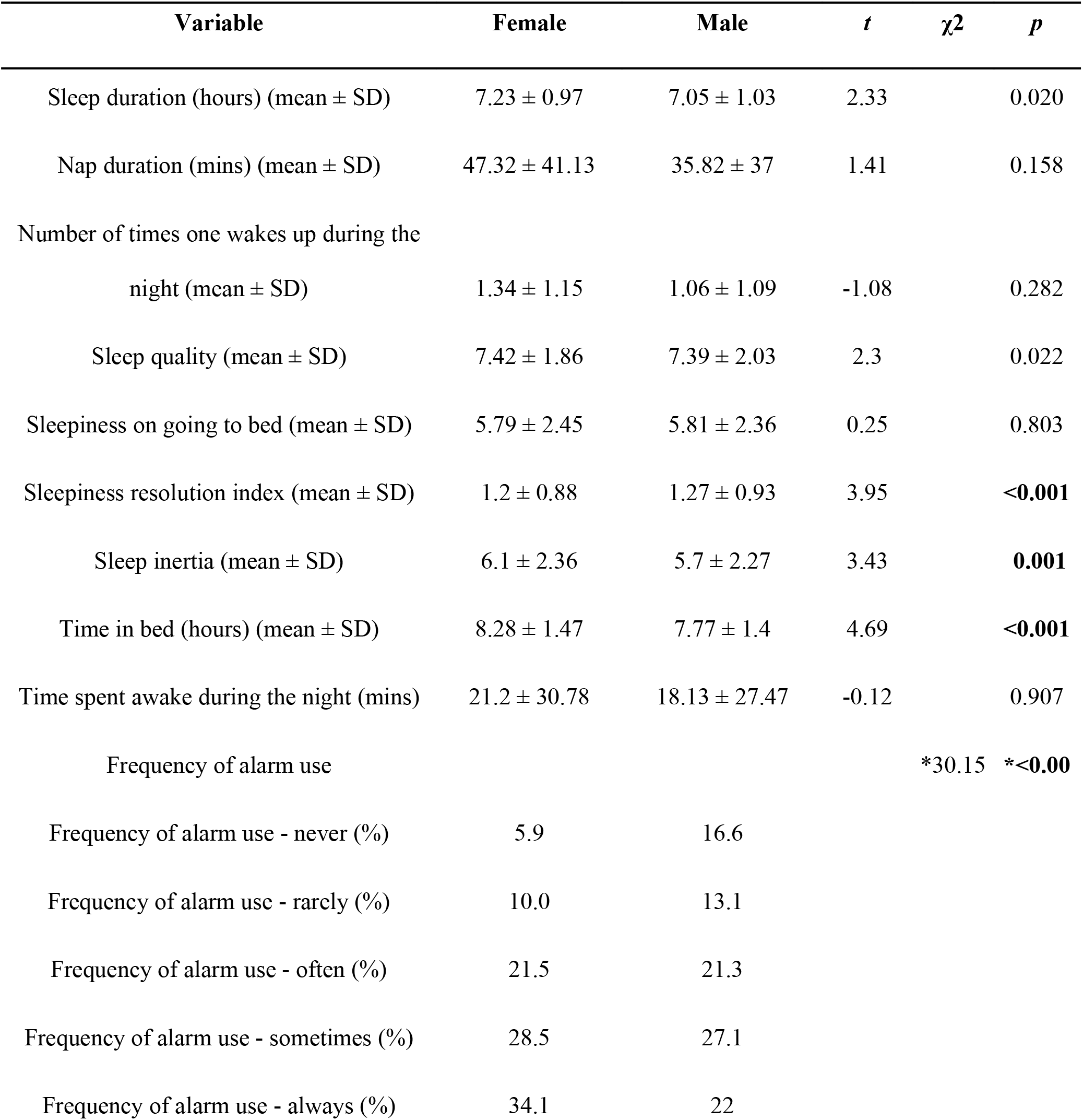
Gender differences in sleep-related variables. Significant P-values are highlighted in bold when applying a bonferroni-corrected alpha threshold of 0.005. *For the categorical variable of frequency of alarm use, a single effect size and p-value are printed given that each of the categories were compared with one another.

| Variable | Female | Male | *t* | $\chi^2$ | *p* |
| --- | --- | --- | --- | --- | --- |
| Sleep duration (hours) (mean ± SD) | 7.23 ± 0.97 | 7.05 ± 1.03 | 2.33 | | 0.020 |
| Nap duration (mins) (mean ± SD) | 47.32 ± 41.13 | 35.82 ± 37 | 1.41 | | 0.158 |
| Number of times one wakes up during the<br>night (mean ± SD) | 1.34 ± 1.15 | 1.06 ± 1.09 | -1.08 | | 0.282 |
| Sleep quality (mean ± SD) | 7.42 ± 1.86 | 7.39 ± 2.03 | 2.3 | | 0.022 |
| Sleepiness on going to bed (mean ± SD) | 5.79 ± 2.45 | 5.81 ± 2.36 | 0.25 | | 0.803 |
| Sleepiness resolution index (mean ± SD) | 1.2 ± 0.88 | 1.27 ± 0.93 | 3.95 | | **<0.001** |
| Sleep inertia (mean ± SD) | 6.1 ± 2.36 | 5.7 ± 2.27 | 3.43 | | **0.001** |
| Time in bed (hours) (mean ± SD) | 8.28 ± 1.47 | 7.77 ± 1.4 | 4.69 | | **<0.001** |
| Time spent awake during the night (mins) | 21.2 ± 30.78 | 18.13 ± 27.47 | -0.12 | | 0.907 |
| Frequency of alarm use |  |  |  | *30.15 | **\*<0.00** |
| Frequency of alarm use - never (%) | 5.9 | 16.6 |  |  |  |
| Frequency of alarm use - rarely (%) | 10.0 | 13.1 |  |  |  |
| Frequency of alarm use - often (%) | 21.5 | 21.3 |  |  |  |
| Frequency of alarm use - sometimes (%) | 28.5 | 27.1 |  |  |  |
| Frequency of alarm use - always (%) | 34.1 | 22 |  |  |  |

### Multivariate analysis - in males, shorter reported sleep was associated with worse navigation

Based on an initial bivariate analysis, difficulty waking up, sleep inertia, sleepiness resolution index, time spent awake during the night and sleep quality were chosen as sleep variables of interest to include in our multivariate linear regression model, given that these variables had the strongest associations with weighted wayfinding distance (see Supplementary Table S3 for the bivariate analysis results). Interactions between gender and each of the sleep variables were included in the model. As for the partial correlations, age, gender, highest level of education achieved, hours of weekly video gaming, hours of weekly phone use, daily hours of sunlight, BMI, weekly units of alcohol, daily cups of caffeine, smoking frequency and frequency of daily strenuous physical activity were included as covariates in the model (Table 3). Before running the regression, as a check for multicollinearity, we calculated the variance inflation factor (*VIF*) for each predictor variable in the model. This revealed that all the predictor variables had a *VIF* value that was less than 5, indicating that multicollinearity was not a concern (Supplementary Table S1). Please see Figure 2 for correlations between each of the sleep variables included in the model.

**Figure 2.**
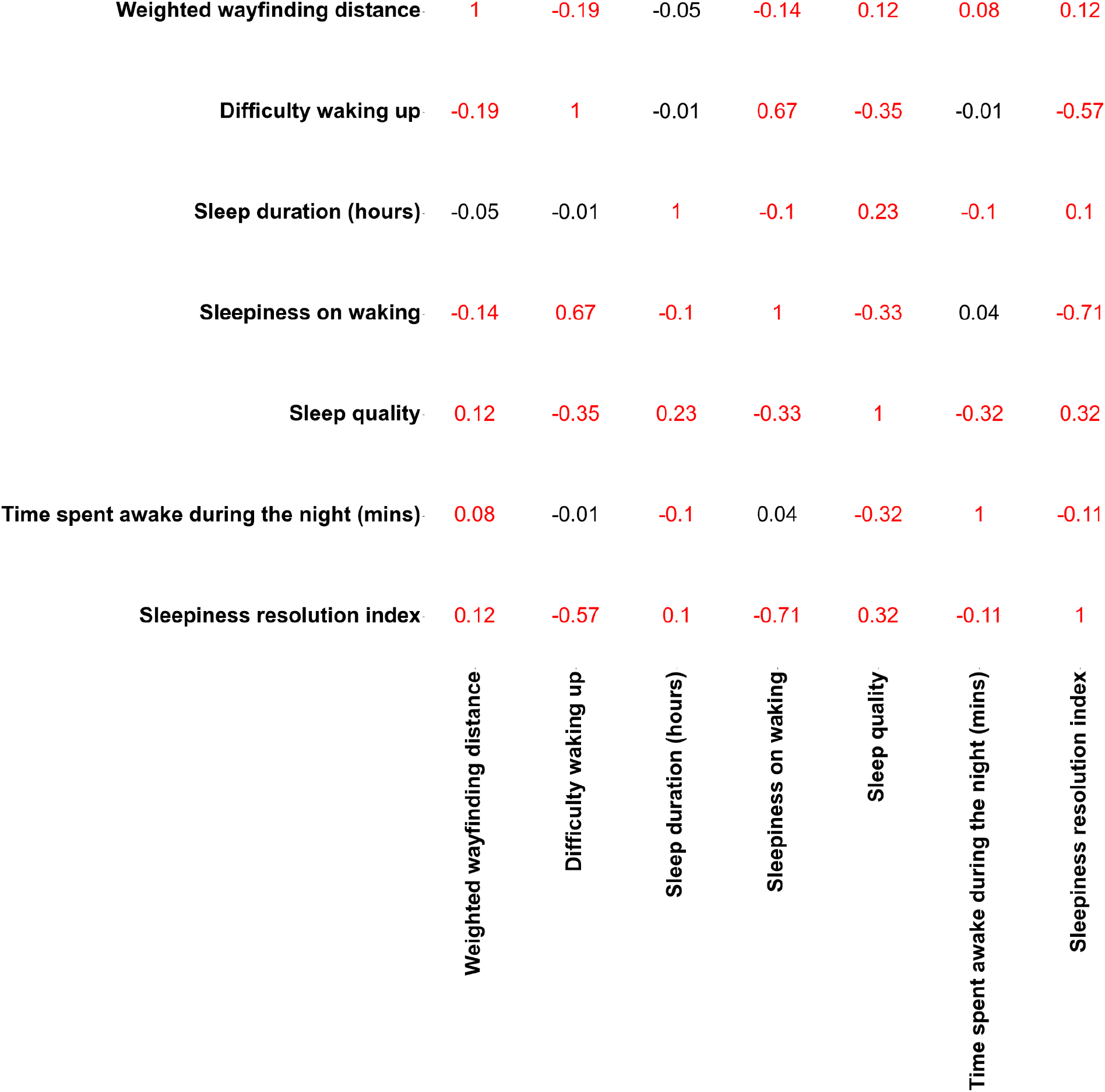
Associations between each of the sleep variables included in the final model and wayfinding distance. Spearman’s correlation coefficients for the associations between each of the sleep variables (sleep quality, sleep inertia, sleepiness resolution index, sleep duration (hours), time spent awake during the night and difficulty waking up) included in the model and and wayfinding distance (across game levels across participants). Values highlighted in red represent significant associations (*p* < 0.05).

### Model

Weighted wayfinding distance ∼ Sleep duration + Sleep duration*Gender + Difficulty waking up + Difficulty waking up*Gender + Sleepiness resolution index + Sleepiness resolution index*Gender + Sleep quality + Sleep quality*Gender + Time spent awake during the night + Time spent awake during the night*Gender + Sleep inertia + Sleep inertia*Gender + Gender + Age + Weekly hours of video gaming + Weekly hours of phone use + BMI + Daily hours of sunlight + Highest education level achieved + Daily cups of caffeine + Weekly units of alcohol + Frequency of daily strenuous physical activity + Smoking frequency

### Sleep-related variables

There was a significant interaction between sleep duration and gender, with longer sleep being associated with shorter wayfinding distance and therefore better navigation performance in men (*β* = -0.11, *f2* = 0.01, *p* = 0.032, *CI* = [-0.22, -0.01])(Table 4 and Figure 3). Bonferroni-corrected post-hoc t-tests showed that men who slept for more than 7 hours were significantly better at navigating than women (*t* = 3.56, *p* < 0.001). However, in those who slept for 7 hours or less, this gender difference in navigation ability was no longer significant (*t* = 1.31, *p* = 0.191)(Table 5). There were no significant associations between sleep quality, difficulty waking up, sleep inertia, sleepiness resolution index or time spent awake during the night and wayfinding distance (*p* > 0.05)(Table 4 and Figure 3).

**Figure 3.**
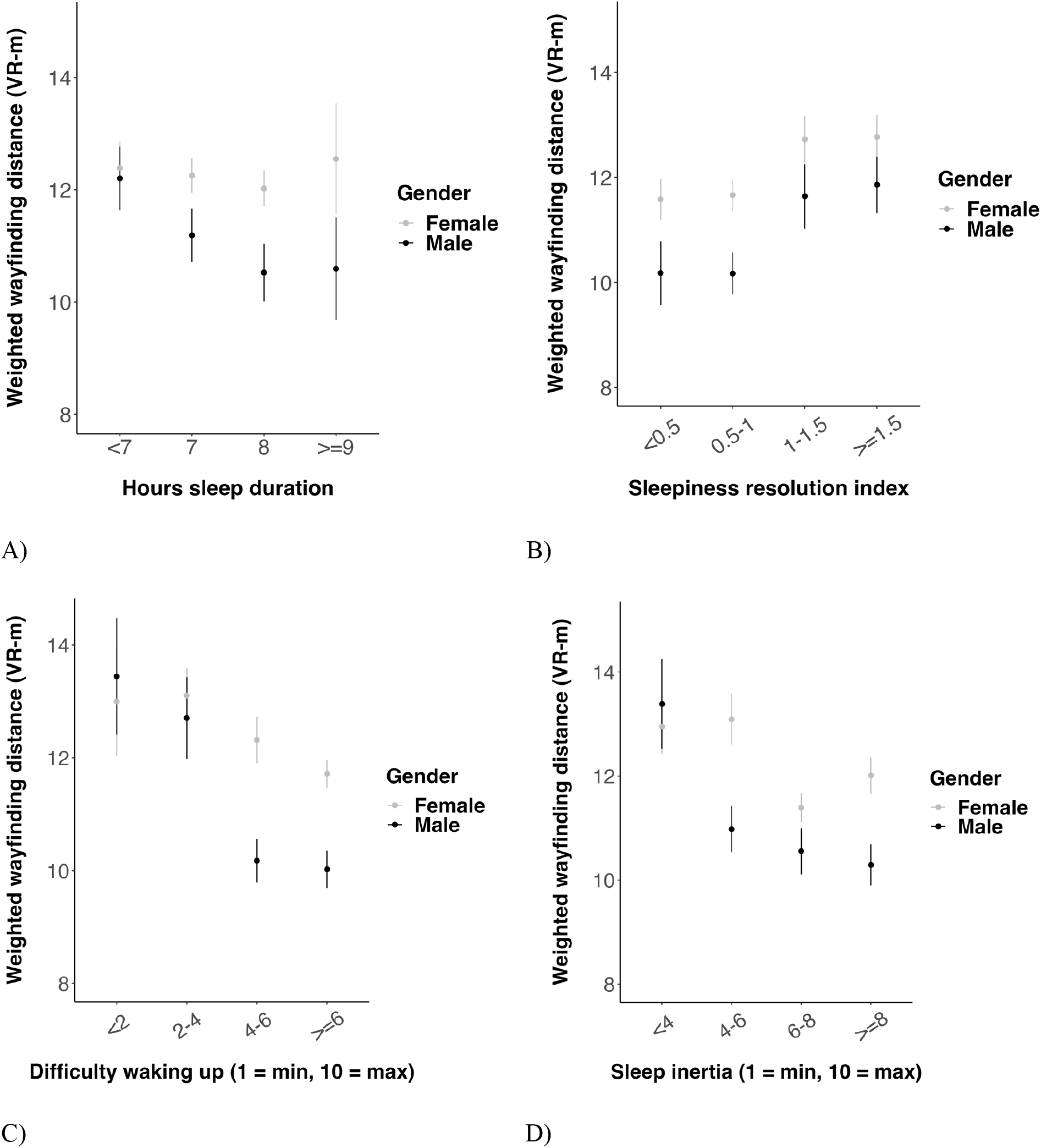
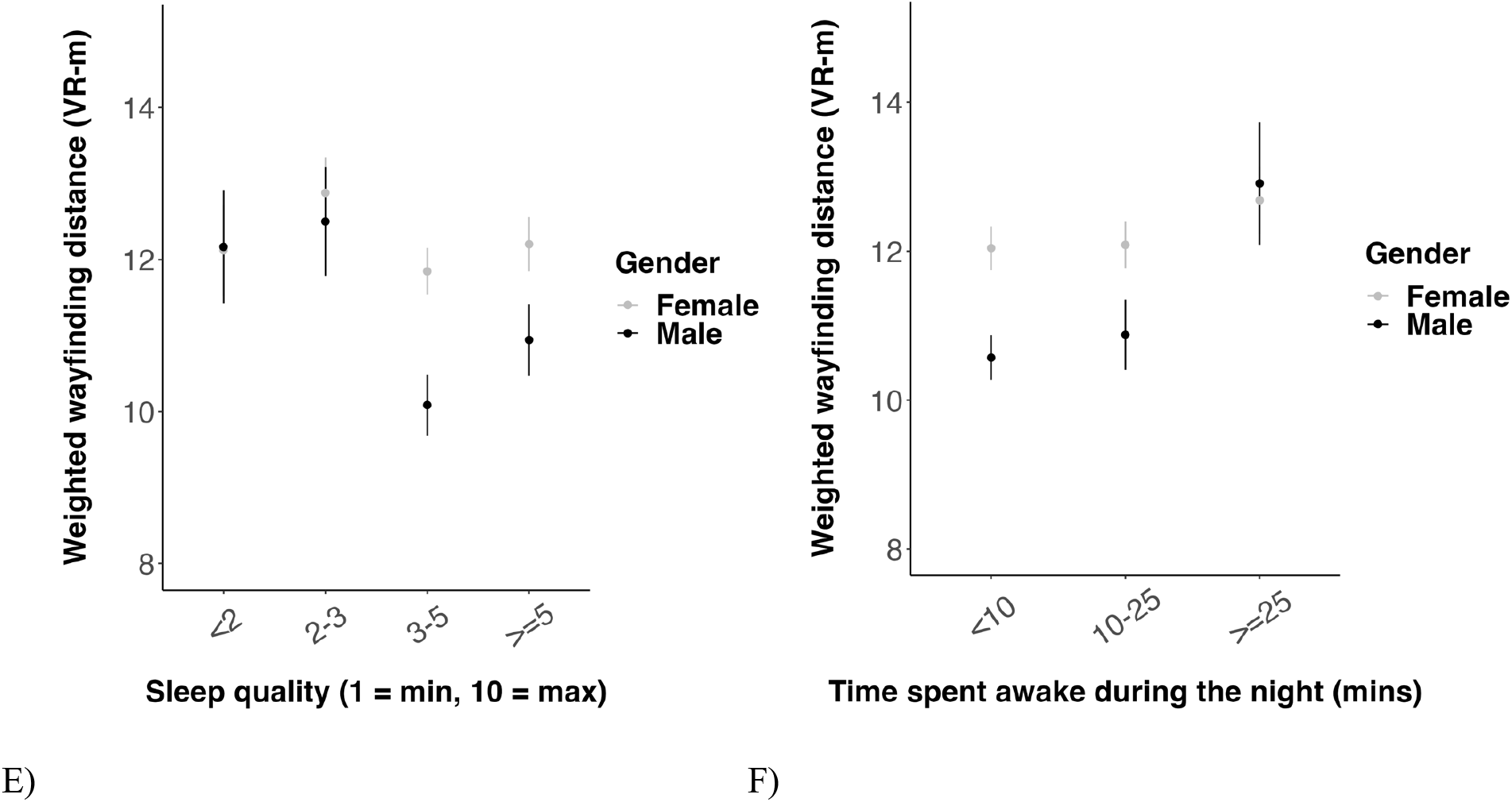
Associations between each of the sleep-related variables and wayfinding distance in men and women. Higher wayfinding distance indicates worse navigation ability. VR-m = Virtual reality meters. (A) The asterix indicates the significant interaction between gender and sleep duration at 5 hours of sleep duration (* = p < 0.05). (B) A value of 1 indicates that the level of sleepiness when going to bed = the level of sleepiness when waking up, a value exceeding 1 indicates that the level of sleepiness on going to bed > the level of sleepiness when waking up and a value of less than 1 indicates that the level of sleepiness on going to bed < level of sleepiness when waking waking up. (C) 1 = minimum difficulty waking up, 10 = maximum difficulty waking up. (D) 1 = minimum sleep inertia, 10 = maximum sleep inertia. (E) 1 = minimum sleep quality, 10 = maximum sleep quality. (A-F) Data points represent the mean wayfinding distance across game levels across participants. Bars represent one standard error above and below the mean wayfinding distance across game levels across participants.

**Table 3.** Model specification with sleep duration as a linear term. The outputs from the multivariate linear regression model are as follows:

**Table 4.**
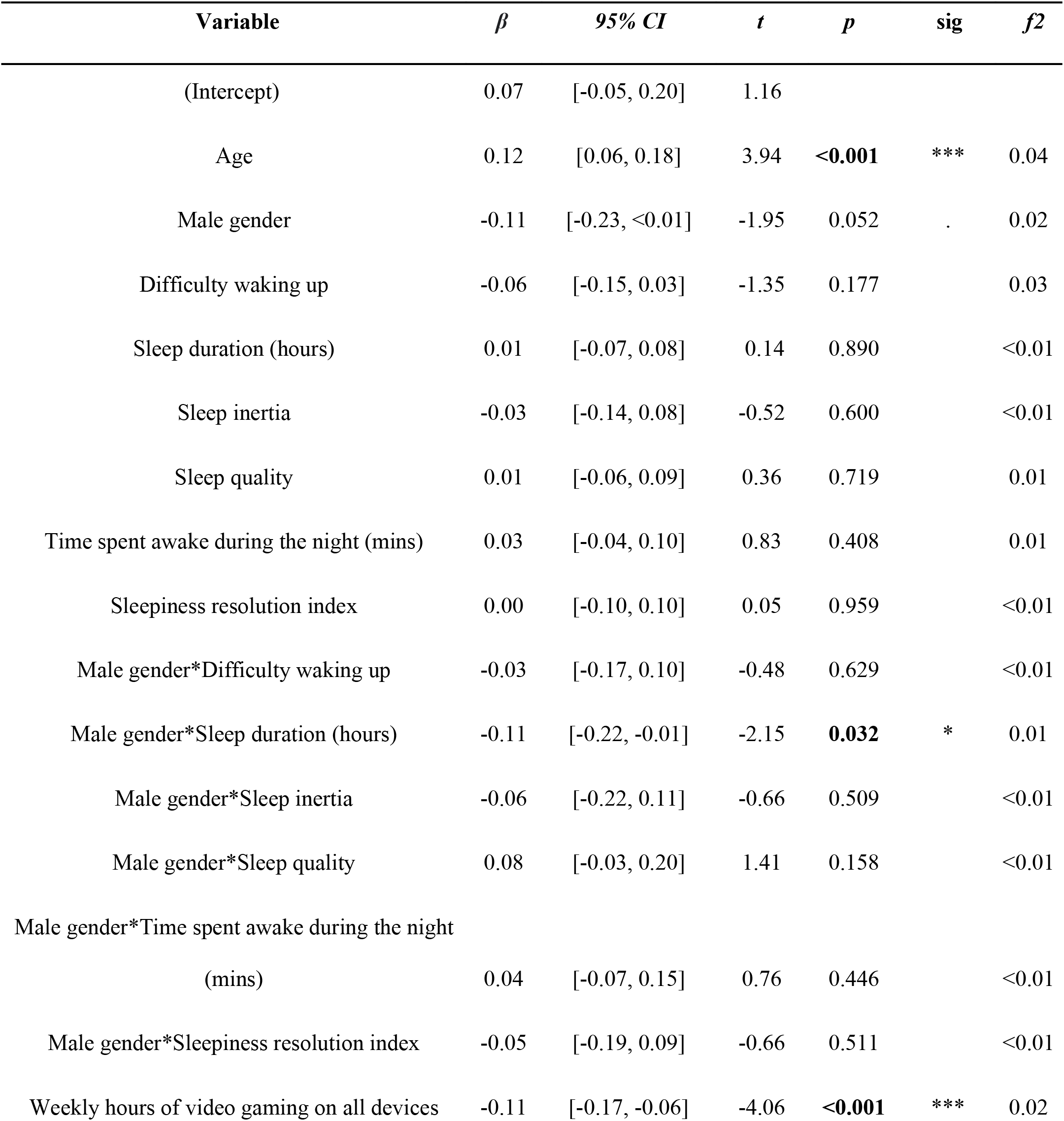
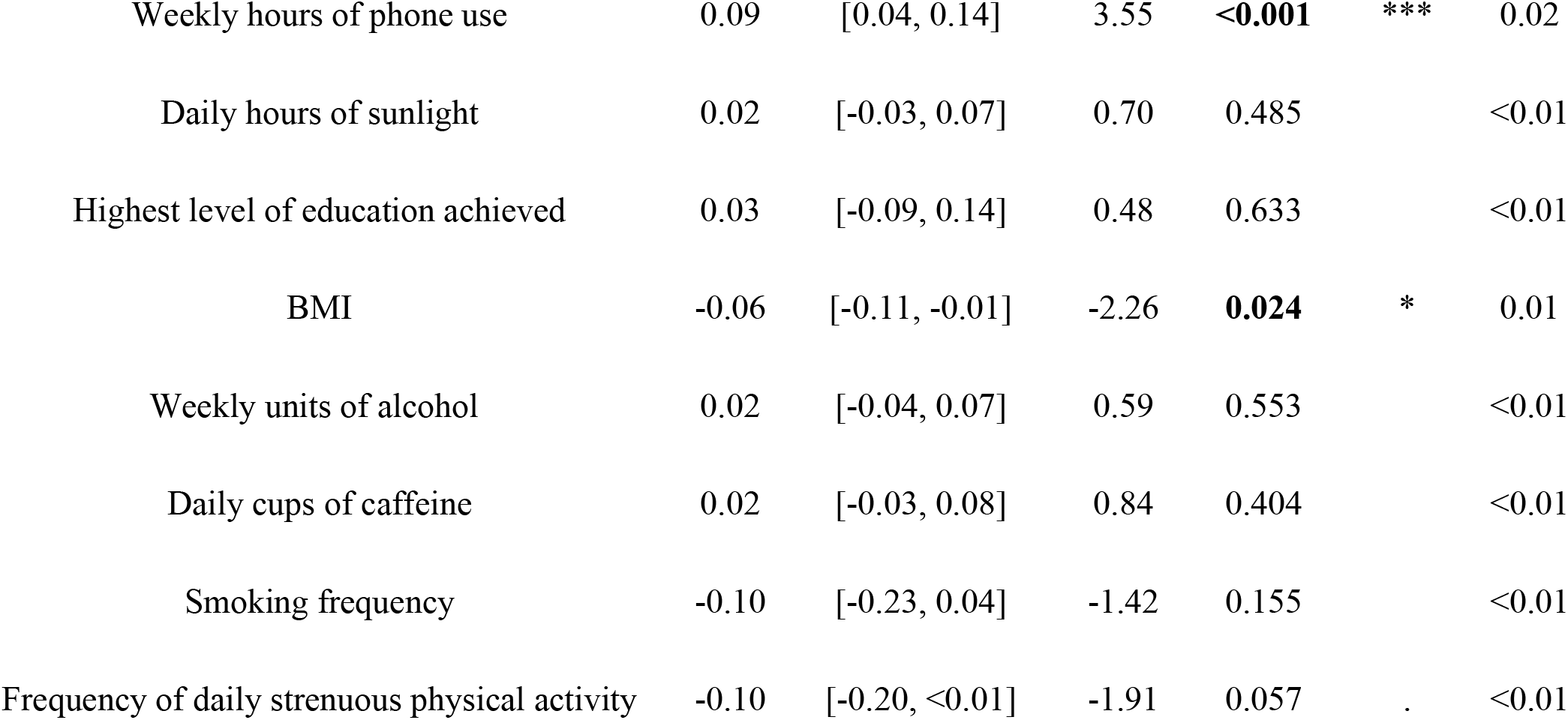
Results of the multiple linear regression model associating sleep-related variables with wayfinding distance when controlling for associated covariates. P-values for the significant associations are highlighted in bold. See this recent study^39^ for consideration of the impact of video games and phone use variables.

| Variable | $\beta$ | 95% CI | $t$ | $p$ | sig | $f^2$ |
| --- | --- | --- | --- | --- | --- | --- |
| (Intercept) | 0.07 | [-0.05, 0.20] | 1.16 |  |  |  |
| Age | 0.12 | [0.06, 0.18] | 3.94 | **<0.001** | *** | 0.04 |
| Male gender | -0.11 | [-0.23, <0.01] | -1.95 | 0.052 | . | 0.02 |
| Difficulty waking up | -0.06 | [-0.15, 0.03] | -1.35 | 0.177 |  | 0.03 |
| Sleep duration (hours) | 0.01 | [-0.07, 0.08] | 0.14 | 0.890 |  | <0.01 |
| Sleep inertia | -0.03 | [-0.14, 0.08] | -0.52 | 0.600 |  | <0.01 |
| Sleep quality | 0.01 | [-0.06, 0.09] | 0.36 | 0.719 |  | 0.01 |
| Time spent awake during the night (mins) | 0.03 | [-0.04, 0.10] | 0.83 | 0.408 |  | 0.01 |
| Sleepiness resolution index | 0.00 | [-0.10, 0.10] | 0.05 | 0.959 |  | <0.01 |
| Male gender*Difficulty waking up | -0.03 | [-0.17, 0.10] | -0.48 | 0.629 |  | <0.01 |
| Male gender*Sleep duration (hours) | -0.11 | [-0.22, -0.01] | -2.15 | **0.032** | * | 0.01 |
| Male gender*Sleep inertia | -0.06 | [-0.22, 0.11] | -0.66 | 0.509 |  | <0.01 |
| Male gender*Sleep quality | 0.08 | [-0.03, 0.20] | 1.41 | 0.158 |  | <0.01 |
| Male gender*Time spent awake during the night<br>(mins) | 0.04 | [-0.07, 0.15] | 0.76 | 0.446 |  | <0.01 |
| Male gender*Sleepiness resolution index | -0.05 | [-0.19, 0.09] | -0.66 | 0.511 |  | <0.01 |
| Weekly hours of video gaming on all devices | -0.11 | [-0.17, -0.06] | -4.06 | **<0.001** | *** | 0.02 |
| Weekly hours of phone use | 0.09 | [0.04, 0.14] | 3.55 | **<0.001** | *** | 0.02 |
| Daily hours of sunlight | 0.02 | [-0.03, 0.07] | 0.70 | 0.485 |  | <0.01 |
| Highest level of education achieved | 0.03 | [-0.09, 0.14] | 0.48 | 0.633 |  | <0.01 |
| BMI | -0.06 | [-0.11, -0.01] | -2.26 | **0.024** | * | 0.01 |
| Weekly units of alcohol | 0.02 | [-0.04, 0.07] | 0.59 | 0.553 |  | <0.01 |
| Daily cups of caffeine | 0.02 | [-0.03, 0.08] | 0.84 | 0.404 |  | <0.01 |
| Smoking frequency | -0.10 | [-0.23, 0.04] | -1.42 | 0.155 |  | <0.01 |
| Frequency of daily strenuous physical activity | -0.10 | [-0.20, <0.01] | -1.91 | 0.057 | . | <0.01 |

**Table 5.**
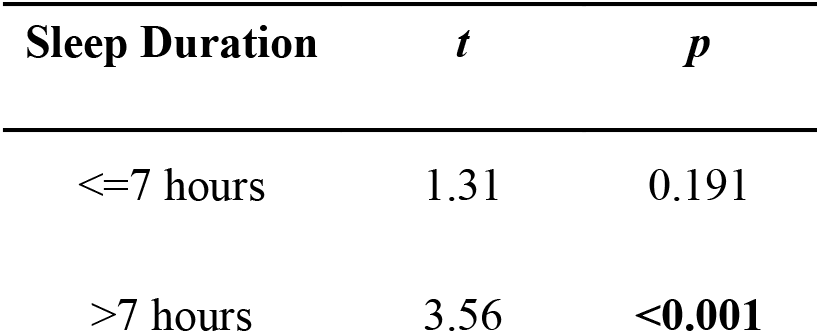
Bonferroni-corrected post-hoc t-tests comparing wayfinding distance in those who reported sleeping <=7 hours with those who slept >7 hours on a typical night. P-values for the significant associations are highlighted in bold when applying a bonferroni-corrected alpha threshold of 0.025.

#### Complementary analysis

### Conducting the model with sleep duration as a quadratic term

When conducting the model with sleep duration specified as a quadratic term (Table 6), there was no significant interaction between sleep duration and gender (*β* = 0.04, *f2* = <0.01, *p* = 0.251, *CI* = [-0.03, 0.12])(Table 7).

**Table 6.**
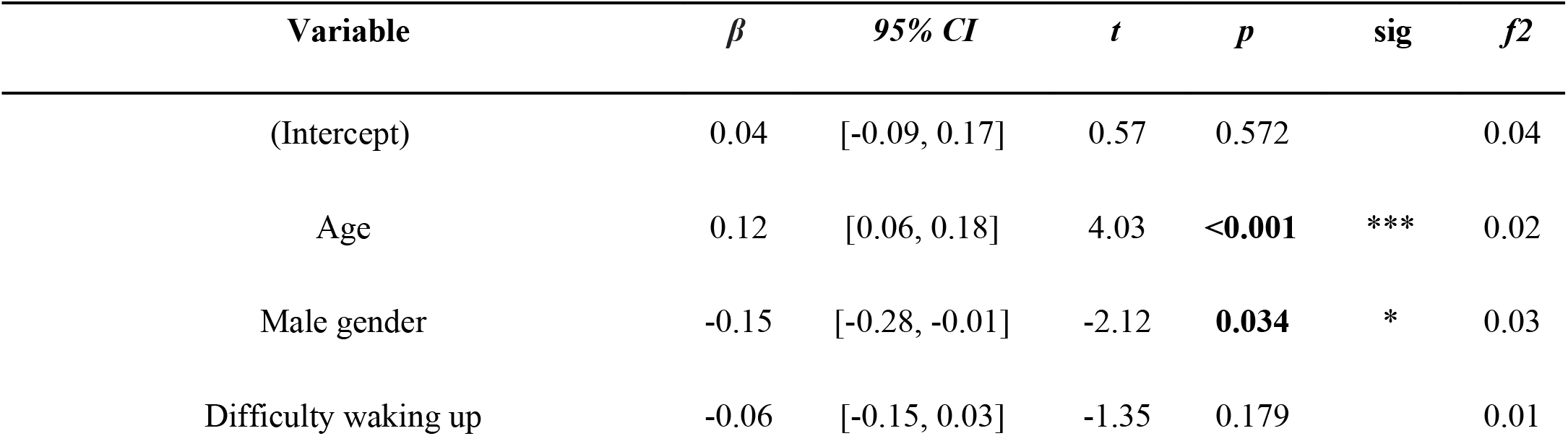
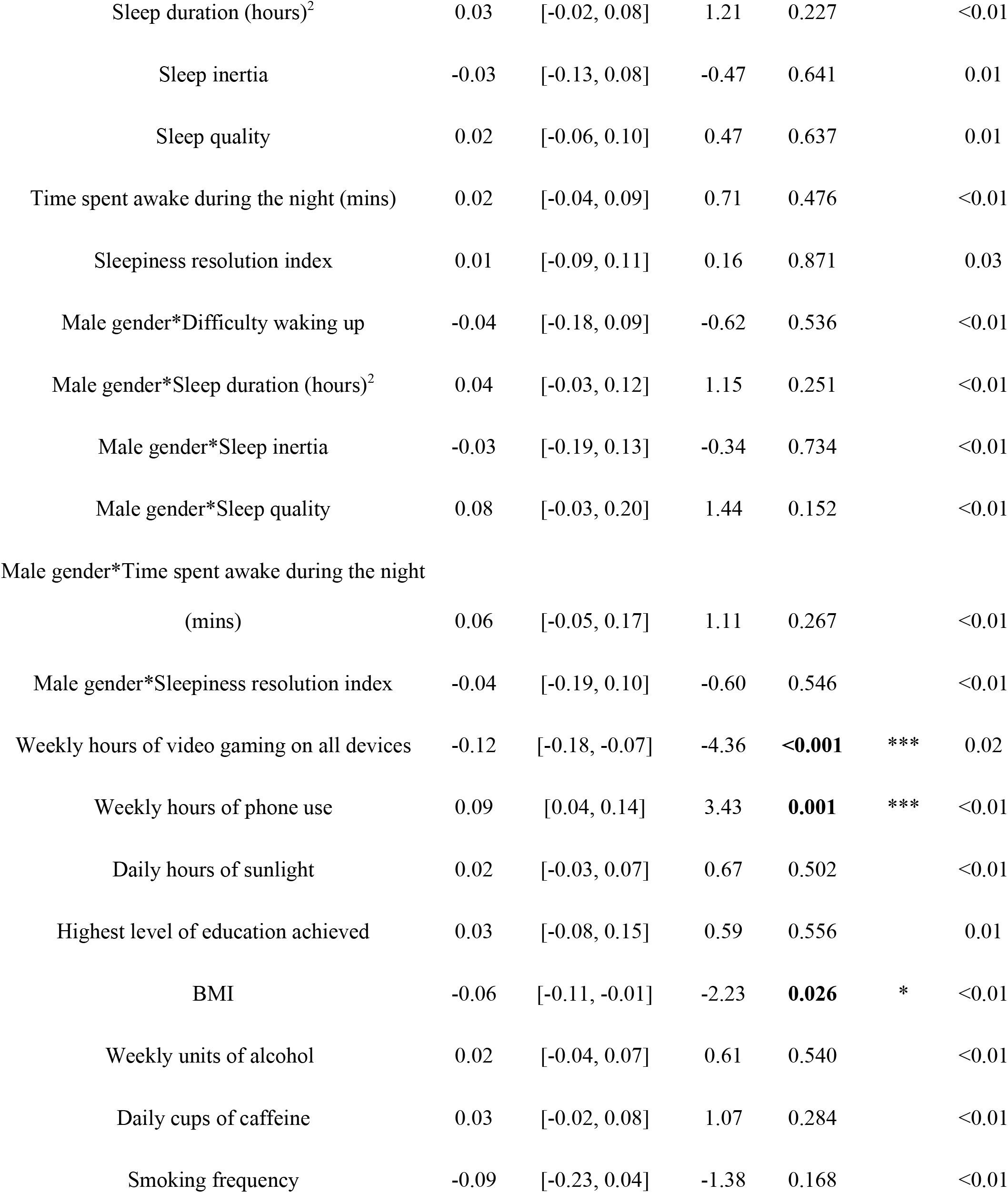
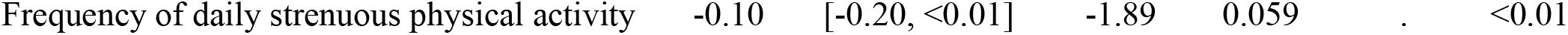
Model specification with sleep duration as a quadratic term.

**Table 7.** Results of the multiple regression model associating sleep-related variables with wayfinding distance, specifying sleep duration as a quadratic term, when controlling for associated covariates. P-values for the significant associations are highlighted in bold.

### Model

Weighted wayfinding distance ∼ Sleep duration^2^ + Sleep duration^2^*Gender + Difficulty waking up + Difficulty waking up*Gender + Sleepiness resolution index + Sleepiness resolution index*Gender + Sleep quality + Sleep quality*Gender + Time spent awake during the night + Time spent awake during the night*Gender + Sleep inertia + Sleep inertia*Gender + Gender + Age + Weekly hours of video gaming + Weekly hours of phone use + BMI + Daily hours of sunlight + Highest education level achieved + Daily cups of caffeine + Weekly units of alcohol + Frequency of daily strenuous physical activity + Smoking frequency

### Conducting the linear model with sleep duration as a linear term in a smaller subsample excluding those spending <5 and > 10 hours in bed (n = 658)

We also ran the same model on a smaller subsample of 658 participants after excluding those who slept for >10 hours or <5 hours (rather than >12 or <4 hours in the current sample), given that our recent study associating sleep duration with navigation ability also used this exclusion criteria^16^. The interaction between sleep duration and gender in this model was not significant (*β **=*** 0.08, *f2* = <0.01, *p* = 0.167, *CI* = [-0.03, 0.19])(Supplementary Table S6). However, sleep duration (hours) showed a significant negative association with wayfinding distance (*β **=*** -0.10, *f2* = <0.01, *p* = 0.018, *CI* = [-0.18, - 0.02])(Supplementary Table S6), where greater hours of sleep were associated with a shorter wayfinding distance and thus better navigation performance. Additionally, the time spent awake during the night (minutes) showed a significant positive association with wayfinding distance (*β **=*** 0.09, *f2* = 0.01, *p* = 0.045, *CI* = [<0.01, 0.17])(Supplementary Table S6), where a greater time spent awake during the night was associated with a longer wayfinding distance and thus worse navigation ability.

## Results Summary

Taken together, there are 3 key findings: 1) greater self-reported sleep duration was associated with significantly better navigation performance only in men when specifying sleep duration as a linear term, 2) self-reported sleep duration did not show a significant interaction with gender when specifying sleep duration as a quadratic term, 3) neither self-reported difficulty waking up, sleep inertia, sleepiness resolution index, sleep quality or time spent awake during the night were significant independent predictors of navigation performance. We also found that when restricting the sleep range to match a previous analysis that only included self-reported sleep duration, greater self-reported difficulty waking up and a shorter self-reported sleep duration .

## Discussion

Here, we investigated the association between multiple sleep-related variables and human spatial navigation performance. Our study is novel as it is the first to investigate the independent association between various sleep-related variables and human spatial navigation performance when accounting for other sleep-related variables.

We found that men reporting 7 hours or less sleep on a typical night had worse wayfinding performance. This supports our hypothesis that sleep duration would show a significant linear association with navigation ability that would differ by gender. This finding aligns with past findings of a gender difference in the effect of sleep duration on cognitive function assessed using measures of verbal fluency, memory and time orientation in a large cohort^25^. It also aligns with more recent findings of those sleeping 6 hours or less on a typical night having worse general cognitive function^43^, although this latter study did not investigate how this association may differ by gender. Our finding of men reporting 7 hours or less sleep having worse wayfinding performance also adds to the previous literature by demonstrating that potentially negative effects of shorter sleep duration may extend from non-spatial to spatial domains of cognitive function. However, it disagrees with studies where no gender differences have been found in the association between sleep duration and cognitive function in both navigation and non-navigation-related domains^13, 44, 45^. Several factors may have accounted for these discrepancies, such as circadian rhythms and the time one spends awake during the day, which have been found to influence gender differences in cognitive function and which were not examined here^46^. Additionally, daily physical activity has been associated with better sleep duration and quality^35^, which in turn may have been associated with better navigation ability. However, we controlled for the frequency of daily strenuous physical activity in our analysis, suggesting that this explanation is unlikely.

We found no significant linear association between either sleep quality or time spent awake during the night and human spatial navigation performance. These findings may seem to contradict previous findings showing that increased sleep fragmentation is associated with poorer wayfinding performance^9^, and that poorer sleep quality is associated with a reduced ability to use shortcuts during wayfinding and thus less efficient wayfinding performance^8^. However, we did find a significant association between the time spent awake during the night - as well as sleep duration - with navigation ability in our smaller subsample analysis (conducted to match a recent study^16^), perhaps suggesting that these aspects of sleep play a more important in role in shaping one’s navigation ability than others. However, another large cohort study using a range of self-reported sleep parameters found no association between any of these parameters and longitudinal change in episodic memory function^47^, suggesting perhaps that performance only in certain domains of cognitive function may be associated with sleep quality. Nonetheless, differences in our experimental task, study design (cross-sectional vs longitudinal) and samples used in our study compared with former studies may have accounted for discrepancies between our findings and those of previous studies. Further investigation is therefore warranted to firmly establish the directionality of any associations between sleep and navigation ability.

When conducting a model with self-reported sleep duration specified as a quadratic term, we did not find a significant association between self-reported sleep duration and human spatial navigation performance. This finding may contradict previous findings showing that those with mid-range sleep durations have better cognitive performance than those with shorter and longer sleep durations, in both navigation-related^16^ and non-navigation-related cognitive tasks^7, 25, 28, 30, 45, 48, 49^. For example, a recent study found a U-shaped association between sleep duration and wayfinding performance on SHQ, where 7 hours of self-reported sleep on a typical night was associated with optimal wayfinding performance^16^. Several factors may have accounted for the discrepancy between this latter finding and our findings. Our sample was solely US-based, whilst this latter study used participants from all around the world^16^. Age may also be another key factor. Our sample was a relatively young sample (mean age = 26.4 years), whilst this latter study used participants with a mean age of 38.7 years^16^. Additionally, other studies have also shown that participants in the middle range of the cohort in terms of age had better cognitive performance at shorter and longer sleep durations than both younger and older participants^30^. Moreover, a recent study showed that in participants of a similar age to ours, sleep can facilitate performance on memory-based tasks but not spatial navigation performance^15^. Another study found sleep fragmentation to be associated with poorer wayfinding performance only in older participants but not in younger participants^7^. Thus, the relatively young age of our sample may have accounted for discrepancies between previous findings and ours. Nonetheless, younger participants have previously been found to have significantly improved wayfinding performance following a night’s sleep whilst older participants did not^9^. Thus, further investigation in samples of a wider age range is warranted to determine the extent to which the association between sleep and spatial navigation performance may differ across the lifespan.

We did not find a significant interaction between gender and any of the sleep-related variables we measured on human spatial navigation performance. One possible explanation for this could be due to gender differences in other variables such as gaming experience, which we also included in our model, that has previously been shown to be associated with wayfinding performance on SHQ^39^ and was shown to mitigate differences in other domains of spatial cognition including mental rotation and perspective taking^50^. Thus, controlling for video game experience may have mitigated gender differences in the association between self-reported sleep duration, self-reported sleep quality and self-reported time spent awake during the night and human spatial navigation performance. Additionally, a recent study demonstrated that a greater level of stress-induced motivation in women can diminish gender differences in spatial navigation performance on a virtual navigation task^51^. Thus, gender differences in other factors may have also mitigated potential differences in the association between self-reported sleep variables and human spatial navigation performance.

Finally, it is important to note that poorer sleep may not always lead to changes in cognitive function. For example, a recent study demonstrated that greater sleep fragmentation was significantly associated with reduced glucose metabolism but not cognitive function^52^. In certain cases, poorer sleep may in fact be associated with better cognitive function, which would be supported by our finding of men with greater sleep inertia, greater difficulty waking up and a lower sleepiness resolution index trending towards having better navigation performance, albeit non-significant associations. In line with this, converting to Lewy-Body Dementia and Parkinson’s Disease Dementia is associated with improved sleep quality, as indicated by a lower severity of insomnia, and having frequent insomnia symptoms has been found to be associated with better cognitive performance^19, 27^. Thus, poorer sleep may be associated with better navigation performance depending on the variables examined, warranting further studies to confirm this hypothesis.

### Limitations and Future Directions

While our study allowed assessment of over 700 participants on an ecologically valid^6^ and reliable cognitive test of navigation^4^, there are a number limitations to consider that future studies could address. Our study relied on self-report measures of sleep duration, resulting in participants being excluded for inaccurately reporting sleep-related measures, as well potential differences in findings due to a discrepancy between self-reported and objective sleep-related measures^22^. Smaller sample sizes for those who slept for shorter and longer sleep durations, where only 5 participants reported sleeping more than 9 hours on a typical night, may have led to an underrepresentation of the association between sleep and navigation ability. Given the cross-sectional nature of our study, future studies should consider an interventional and longitudinal design^17^, especially considering that gender can predict night-to-night variation in sleep duration and differences in sleep duration on weekdays compared with weekends^53^. Incorporating objective sleep-based measures and neural recordings in such studies will help identify individuals who may be more likely to benefit from sleep-based therapeutic interventions and effectively target those at greater risk of cognitive decline. Additionally, conducting these studies across countries and cultures will help determine the extent to which associations between sleep and navigation performance may differ across different populations, as has been shown for both self-reported sleep duration^53^ and self-reported navigation ability^54^. Such future studies would also benefit from using a broader range of virtual navigation and spatial memory tests, that extend to real-world environments^5, 55–59^. This would be useful to help explore our incidental observation that a higher BMI was associated with worse navigation performance, a finding that mirrors evidence that high BMI is associated with worse episodic memory^60^.

### Conclusion

Overall, our findings suggest that the effects of sleep duration on navigation ability may differ by gender. Accordingly, this research will serve as a platform for future research looking into how actively changing one’s sleep habits may improve their navigation ability and provide a greater understanding as to why individual differences in navigation ability may exist.

## Acknowledgements

We would like to thank all the participants who volunteered to take part in this research. This research is part of the Sea Hero Quest initiative funded and supported by Deutsche Telekom. Alzheimer’s Research UK (ARUK-DT2016-1) funded the analysis; Glitchers designed and produced the game; and Saatchi and Saatchi London managed its creation. EY was funded by The Leverhulme Trust. We thank Masoud Tasmanian for helpful comments on this manuscript.

## Data availability

A dataset containing the preprocessed trajectory lengths and demographic information is available at https://osf.io/d5q4r/. We also set up a portal where researchers can invite a targeted group of participants to play SHQ and generate data about their spatial navigation capabilities: https://seaheroquest.alzheimersresearchuk.org/. Those invited to play the game will be sent a unique participant key, generated by the SHQ system according to the criteria and requirements of a specific project decided by the experimenter. Access to the portal will be granted for non-commercial purposes. Future publications based on this dataset should add ‘Sea Hero Quest Project’ as a co-author.

## Code availability

The code used to produce this data is accessible at: https://osf.io/d5q4r/

