## Supplementary Materials for "Shorter self-reported sleep duration in men is associated with worse virtual spatial navigation performance"

| Variable | *VIF* |
|---|---|
| Age | 1.52 |
| Male gender | 1.37 |
| Difficulty waking up | 3.33 |
| Sleep duration (hours) | 2.13 |
| Sleep inertia | 4.91 |
| Sleep quality | 2.64 |
| Time spent awake during the night (mins) | 1.93 |
| Sleepiness resolution index | 4.21 |
| Male gender*Difficulty waking up | 3.31 |
| Male gender*Sleep duration (hours) | 2.08 |
| Male gender*Sleep inertia | 4.64 |
| Male gender*Sleep quality | 2.58 |
| Male gender*Time spent awake during the night (mins) | 1.83 |
| Male gender*Sleepiness resolution index | 3.89 |
| Weekly hours of video gaming on all devices | 1.29 |
| Weekly hours of phone use | 1.11 |
| Daily hours of sunlight | 1.12 |

| | |
|---|---|
| Highest level of education achieved | 1.15 |
| BMI | 1.18 |
| Weekly units of alcohol | 1.14 |
| Daily cups of caffeine | 1.19 |
| Smoking frequency | 1.23 |
| Frequency of daily strenuous physical activity | 1.07 |

**Table S1. Variance inflation factor (*VIF*) values for each of the predictor variables included in the main model when specifying sleep duration as a linear term.**

| **Variable** | ***VIF*** |
|---|---|
| Age | 1.51 |
| Male gender | 1.89 |
| Difficulty waking up | 3.30 |
| Sleep duration (hours) | 1.89 |
| Sleep inertia | 4.93 |
| Sleep quality | 2.52 |
| Time spent awake during the night (mins) | 1.93 |
| Sleepiness resolution index | 4.22 |
| Male gender*Difficulty waking up | 3.28 |

| | |
|---|---|
| Male gender*Sleep duration (hours) | 2.51 |
| Male gender*Sleep inertia | 4.58 |
| Male gender*Sleep quality | 2.55 |
| Male gender*Time spent awake during the night (mins) | 1.82 |
| Male gender*Sleepiness resolution index | 3.87 |
| Weekly hours of video gaming on all devices | 1.29 |
| Weekly hours of phone use | 1.11 |
| Daily hours of sunlight | 1.12 |
| Highest level of education achieved | 1.15 |
| BMI | 1.17 |
| Weekly units of alcohol | 1.14 |
| Daily cups of caffeine | 1.20 |
| Smoking frequency | 1.23 |
| Frequency of daily strenuous physical activity | 1.07 |

**Table S2. Variance inflation factor (*VIF*) values for each of the predictor variables included in the main model when specifying sleep duration as a quadratic term.**

| **Variable** | ***r*** | ***p*** | ***95% CI*** |
|---|---|---|---|
| Difficulty waking up | -0.14 | **<0.001** | [-0.25, -0.04] |

| | | | |
|---|---|---|---|
| Sleepiness resolution index | 0.10 | 0.010 | [-0.01, 0.20] |
| Sleep inertia | -0.09 | 0.015 | [-0.20, 0.02] |
| Sleep quality | 0.06 | 0.088 | [-0.04, 0.17] |
| Time spent awake during the night (mins) | 0.06 | 0.089 | [-0.04, 0.17] |
| Frequency of alarm use | 0.06 | 0.127 | [-0.16, 0.05] |
| Number of times one wakes up during the night | -0.06 | 0.130 | [-0.05, 0.16] |
| Sleep duration (hours) | -0.05 | 0.161 | [-0.16, 0.05] |
| Frequency of naps | 0.04 | 0.235 | [-0.06, 0.15] |
| Sleepiness on going to bed | 0.04 | 0.285 | [-0.07, 0.15] |
| Time in bed (hours) | -0.04 | 0.322 | [-0.14, 0.07] |
| Nap duration (mins) | -0.04 | 0.348 | [-0.14, 0.07] |

**Table S3. Partial spearman's correlations showing the associations between sleep-related variables and weighted wayfinding distance when controlling for age, gender, highest level of education achieved, weekly hours of video gaming, and weekly hours of phone use, BMI, daily hours of sunlight, daily cups of coffee, frequency of daily strenuous physical activity, smoking frequency and weekly units of alcohol as covariates.** P-values highlighted in bold indicate significant associations when applying an alpha threshold of 0.004, bonferroni-corrected with 12 comparisons. *

| **Model** |
|---|

Weighted wayfinding distance ~ Sleep duration + Sleep duration*Gender + Difficulty waking up + Difficulty waking up*Gender + Sleepiness resolution index + Sleepiness resolution index*Gender + Sleep quality + Sleep quality*Gender + Time spent awake during the night + Time spent awake during the night*Gender + Sleep inertia + Sleep inertia*Gender + Gender + Age + Weekly hours of video gaming + Weekly hours of phone use + BMI + Daily hours of sunlight + Highest education level achieved + Daily cups of caffeine + Weekly units of alcohol + Frequency of daily strenuous physical activity + Smoking frequency

**Table S4. Model specification for the model predicted weighted wayfinding distance using sleep-related variables including a subsample of participants who reported spending >= 5 and <= 10 hours in bed.**

| **Variable** | ***VIF*** |
|---|---|
| Age | 1.57 |
| Male gender | 1.38 |
| Difficulty waking up | 4.71 |
| Sleep duration (hours) | 2.37 |
| Sleep inertia | 6.46 |
| Sleep quality | 2.81 |
| Time spent awake during the night (mins) | 2.76 |
| Sleepiness resolution index | 4.21 |
| Male gender*Difficulty waking up | 4.54 |

| | |
|---|---|
| Male gender*Sleep duration (hours) | 2.39 |
| Male gender*Sleep inertia | 6.55 |
| Male gender*Sleep quality | 2.89 |
| Male gender*Time spent awake during the night (mins) | 2.87 |
| Male gender*Sleepiness resolution index | 4.34 |
| Weekly hours of video gaming on all devices | 1.32 |
| Weekly hours of phone use | 1.12 |
| Daily hours of sunlight | 1.10 |
| Highest level of education achieved | 1.15 |
| BMI | 1.19 |
| Weekly units of alcohol | 1.14 |
| Daily cups of caffeine | 1.22 |
| Smoking frequency | 1.24 |
| Frequency of daily strenuous physical activity | 1.07 |

**Table S5. Variance inflation factors for each of the predictor variables included in the model predicting weighted wayfinding distance using sleep-related variables in those spending >= 5 and <= 10 hours in bed.**

| Variable | *β* | *95% CI* | *t* | *p* | sig | *f2* |
|---|---|---|---|---|---|---|

| | | | | | | |
|---|---|---|---|---|---|---|
| (Intercept) | -0.05 | [-0.19, 0.08] | -0.75 | 0.455 | | |
| Age | 0.10 | [0.04, 0.17] | 3.13 | **0.002** | ** | 0.03 |
| Male gender | 0.13 | [0.01, 0.25] | 2.06 | **0.040** | * | 0.03 |
| Difficulty waking up | -0.10 | [-0.22, 0.01] | -1.78 | 0.075 | . | 0.03 |
| Sleep duration (hours) | -0.10 | [-0.18, -0.02] | -2.37 | **0.018** | * | <0.001 |
| Sleep inertia | -0.07 | [-0.21, 0.06] | -1.10 | 0.272 | | <0.001 |
| Sleep quality | 0.08 | [-0.01, 0.17] | 1.78 | 0.076 | . | <0.001 |
| Time spent awake during the night (mins) | 0.09 | [<0.001, 0.17] | 2.01 | **0.045** | * | 0.01 |
| Sleepiness resolution index | -0.04 | [-0.14, 0.07] | -0.67 | 0.501 | | <0.001 |
| Male gender*Difficulty waking up | 0.04 | [-0.11, 0.19] | 0.52 | 0.604 | | <0.001 |
| Male gender*Sleep duration (hours) | 0.08 | [-0.03, 0.19] | 1.38 | 0.167 | | <0.001 |
| Male gender*Sleep inertia | 0.04 | [-0.13, 0.22] | 0.47 | 0.636 | | <0.001 |
| Male gender*Sleep quality | -0.06 | [-0.19, 0.06] | -1.03 | 0.302 | | <0.001 |
| Male gender*Time spent awake during the night (mins) | -0.06 | [-0.17, 0.06] | -0.95 | 0.344 | | <0.001 |
| Male gender*Sleepiness resolution index | 0.06 | [-0.10, 0.21] | 0.72 | 0.471 | | <0.001 |
| Weekly hours of video gaming on all devices | -0.11 | [-0.17, -0.05] | -3.70 | **<0.001** | *** | 0.02 |
| Weekly hours of phone use | 0.09 | [0.04, 0.15] | 3.28 | **<0.001** | ** | 0.02 |
| Daily hours of sunlight | 0.01 | [-0.05, 0.06] | 0.25 | 0.802 | | <0.001 |
| Highest level of education achieved | 0.03 | [-0.09, 0.15] | 0.48 | 0.633 | | <0.001 |

| | | | | | | |
|---|---|---|---|---|---|---|
| BMI | -0.06 | [-0.12, <0.001] | -2.13 | **0.034** | * | 0.01 |
| Weekly units of alcohol | 0.02 | [-0.03, 0.08] | 0.88 | 0.382 | | <0.001 |
| Daily cups of caffeine | 0.02 | [-0.04, 0.07] | 0.58 | 0.565 | | <0.001 |
| Smoking frequency | -0.08 | [-0.22, 0.06] | -1.10 | 0.273 | | <0.001 |
| Frequency of daily strenuous physical activity | -0.11 | [-0.21, <0.001] | -1.95 | 0.052 | . | 0.01 |

**Table S6. Model output for predicting weighted wayfinding distance using sleep-related variables and associated covariates when using a subsample of participants who reported spending >= 5 and <= 10 hours in bed.** P-values for the significant associations are highlighted in bold.

| Sleep duration (hours) | Female | Male |
|---|---|---|
| <7 | 80 | 90 |
| 7 | 172 | 124 |
| 8 | 129 | 93 |
| >=9 | 28 | 21 |

A)

| Sleepiness resolution index | Female | Male |
|---|---|---|
| <0.5 | 75 | 34 |
| 0.5-1 | 122 | 103 |

| | | |
|---|---|---|
| 1-1.5 | 97 | 91 |
| >=1.5 | 116 | 86 |

B)

| Difficulty waking up | Female | Male |
|---|---|---|
| <2 | 34 | 35 |
| 2-4 | 82 | 73 |
| 4-6 | 76 | 64 |
| >=6 | 218 | 142 |

C)

| Sleep inertia | Female | Male |
|---|---|---|
| <4 | 76 | 75 |
| 4-6 | 83 | 73 |
| 6-8 | 118 | 109 |
| >=8 | 132 | 71 |

D)

| Sleep quality | Female | Male |
|---|---|---|
| <2 | 46 | 42 |
| 2-3 | 96 | 85 |
| 3-5 | 144 | 101 |
| >=5 | 123 | 100 |

E)

| Time spent awake during the night (minutes) | Female | Male |
|---|---|---|
| <10 | 109 | 81 |
| 10-25 | 190 | 164 |
| >=25 | 110 | 83 |

F)

**Tables S7A-F. Number of men and women in each category of sleep duration (hours), sleepiness resolution index, difficulty waking up, sleep inertia, sleep quality and time spent awake during the night (minutes).**

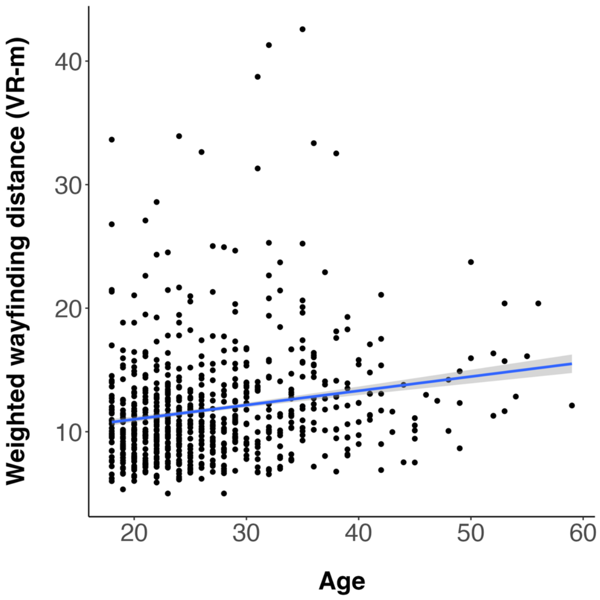

Weighted wayfinding distance (VR-m)
40
30
20
10
20
30
40
50
60
Age


A)

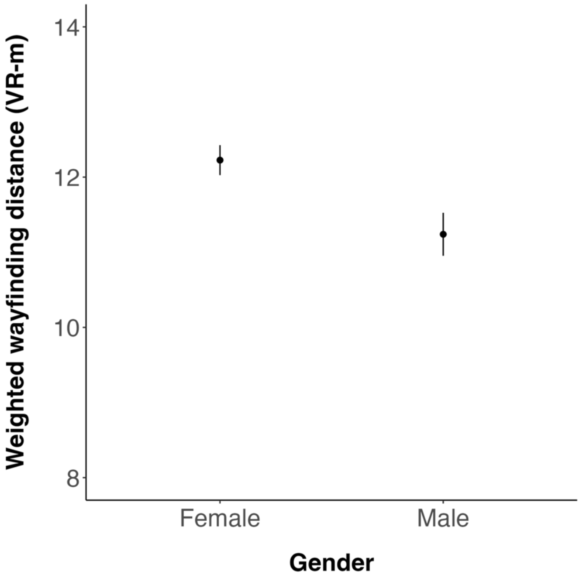

Weighted wayfinding distance (VR-m)
14
12
10
8
Female
Male
Gender


B)

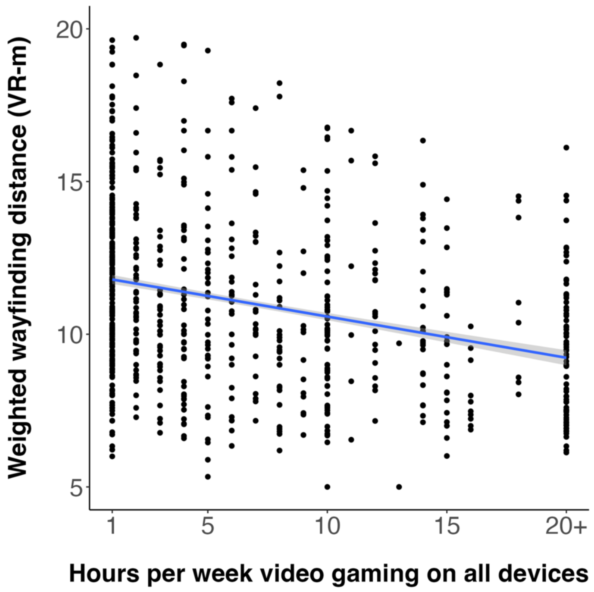
Weighted wayfinding distance (VR-m)
20
15
10
5
1
5
10
15
20+
Hours per week video gaming on all devices

C)

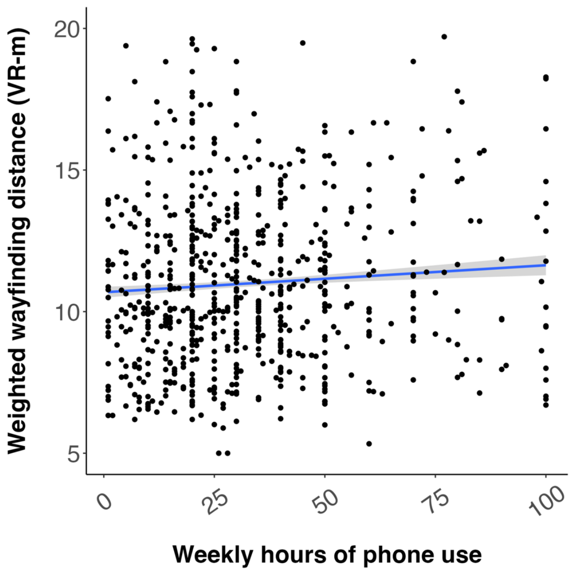
Weighted wayfinding distance (VR-m)
20
15
10
5
0
25
50
75
100
Weekly hours of phone use

D)

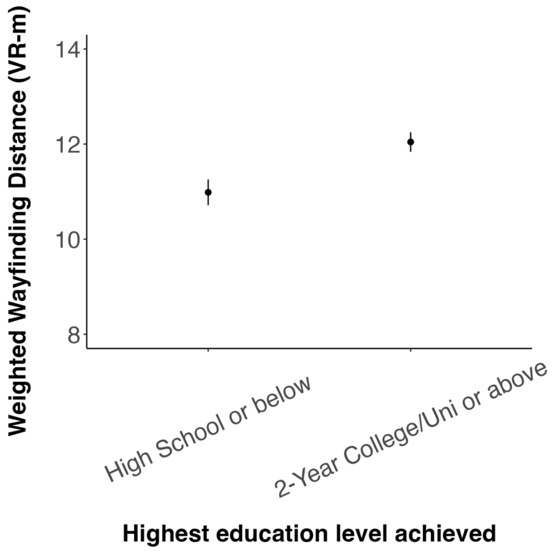
Weighted Wayfinding Distance (VR-m)
14
12
10
8
High School or below
2-Year College/Uni or above
Highest education level achieved

E)

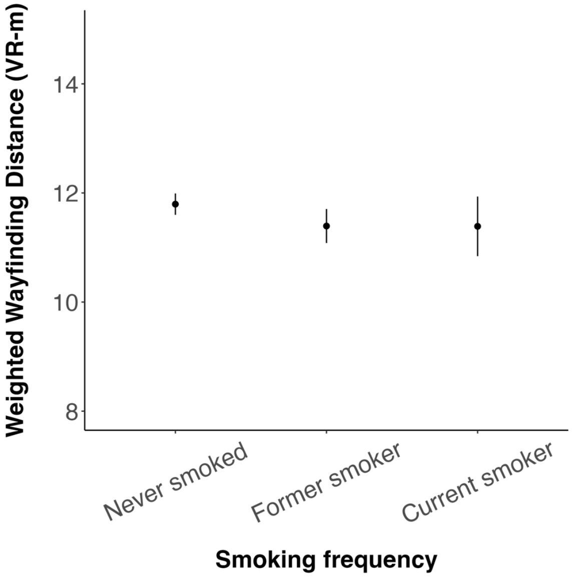
Weighted Wayfinding Distance (VR-m)
14
12
10
8
Never smoked
Former smoker
Current smoker
Smoking frequency

F)

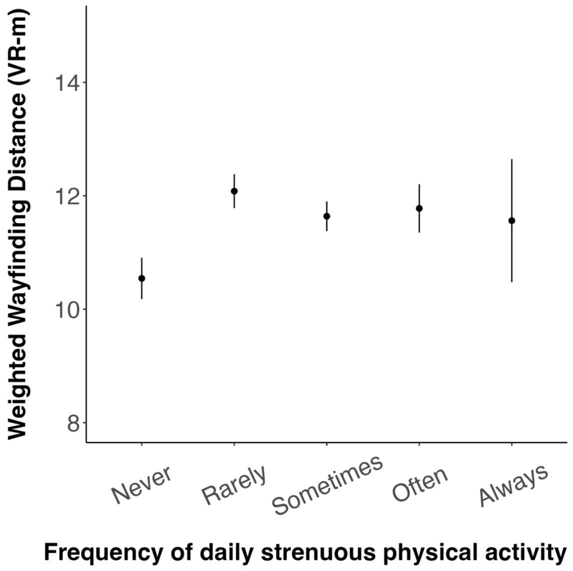
Weighted Wayfinding Distance (VR-m)
14
12
10
8
Never
Rarely
Sometimes
Often
Always
Frequency of daily strenuous physical activity

G)

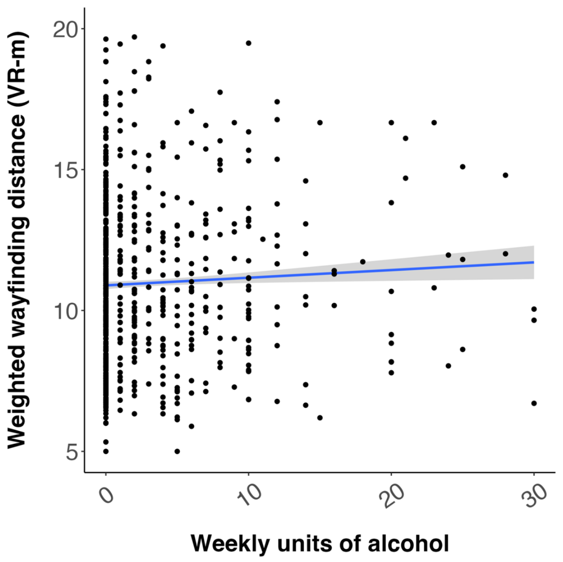
Weighted wayfinding distance (VR-m)
20
15
10
5
0
10
20
30
Weekly units of alcohol

H)

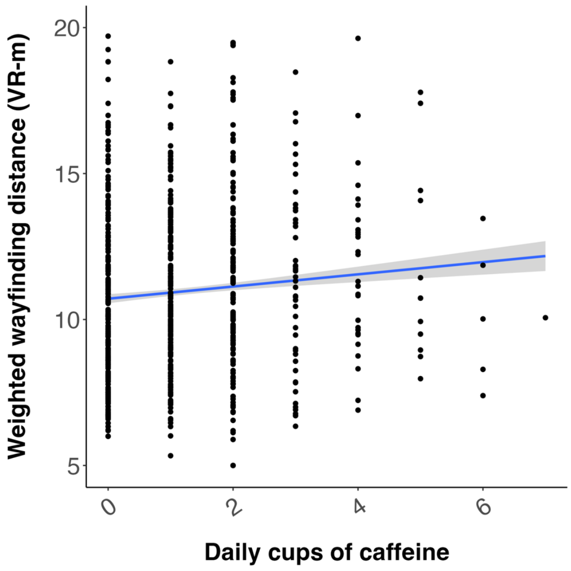
Weighted wayfinding distance (VR-m)
20
15
10
5
0
2
4
6
Daily cups of caffeine

I)

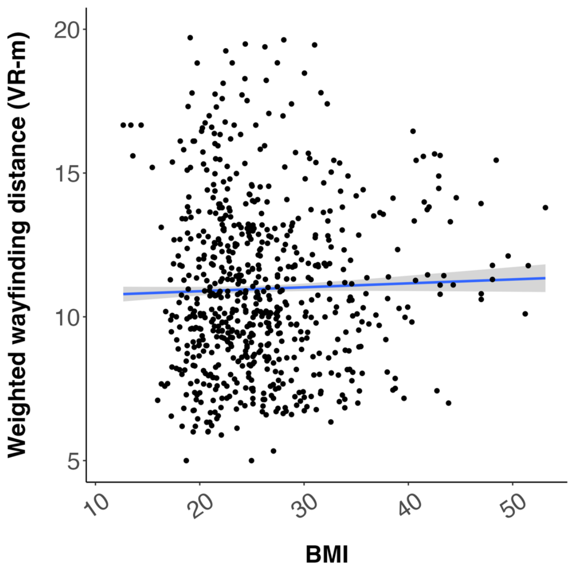
Weighted wayfinding distance (VR-m)
20
15
10
5
10
20
30
40
50
BMI

J)

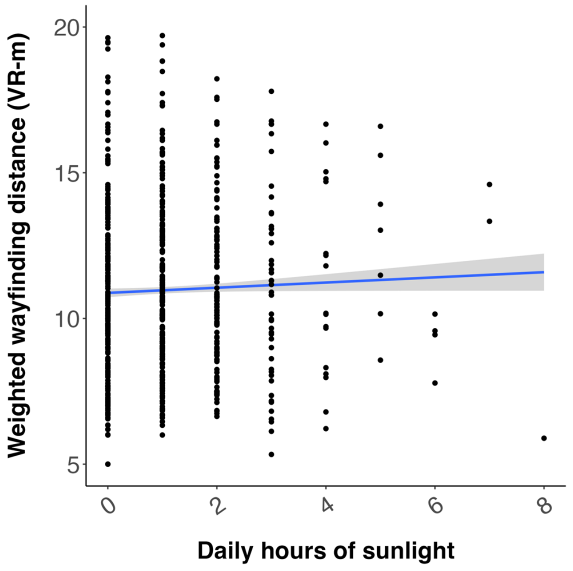


K)

**Figure S1. (A-K) Associations between each of the demographic variables and weighted wayfinding distance.** VR-m = virtual reality metres

(A,C,D,H-K) Blue line indicates the mean wayfinding distance across game levels across participants. Grey shading surrounding the blue line indicates the standard error of the mean corresponding to this wayfinding distance. Data points indicate the mean wayfinding distance across game levels for an individual participant.

(B,E-G) Data points represent the mean wayfinding distance across game levels across participants. Bars represent the standard error of the mean corresponding to this wayfinding distance.

<u>Questionnaires</u>

**<u>Demographics:</u>**

1. How old are you?

2. What gender are you? (male, female, other)

3. What is the highest level of education you have achieved? (no formal education, some formal education, high school, 2-year college/university, 4-year college/university, masters, PhD)

4. Please indicate your height in feet and inches

5. Please enter your weight in pounds*

*Weight was converted to kg in order to calculate Body Mass Index (BMI)

**<u>Gaming experience:</u>**

1. How often do you play video games per week? (0 = minimum, 20+ hours = maximum)

2. How often you do you use a smartphone or a tablet per week? (0 = minimum, 40+ hours = maximum)

**<u>Sleep-related variables:</u>**

1. What time do you go to sleep? (Please enter the time out of 24:00 hours (00 - 12 AM, 12 - 24 PM)

2. How long does it take you to fall asleep? (Please indicate hours on the left and minutes on the right. For example, if it takes an hour and a half to fall asleep, please enter 01 on the left and 30 on the right).

3. How many times do you wake up on a typical night?

4. How long are you awake for, in total, during the night? (please enter your response in minutes; 1 hr = 60, 2hr = 120, 3hr = 180, 4hr = 240)

5. What time do you wake up? Please enter the time out of 24:00 hours (00 - 12 AM, 12 - 24 PM).

6. Does your alarm clock wake you up? (never, rarely, sometimes, often, always)

7. Do you take naps during the day? (never, rarely, sometimes, often, always)

8. If you take naps during the day, how long is the duration of your nap(s) in total? Please enter your response in minutes. Leave the slider at 0 if you do not take naps. (slider, 0 = min, 240 = max)

9. Do you do any strenuous physical activity during the day? (never, rarely, sometimes, often, always)

10. How would you rate your sleep quality overall? (1 = very good, 10 = very bad) *

11. How difficult do you find it to wake up/get up? (1 = very easy, 10 = very hard)

12. Please indicate your level of sleepiness upon waking (0 = extremely alert, 10 = extremely sleepy)**

13. Please indicate your level of sleepiness upon bedtime (0 = extremely alert, 10 = extremely sleepy)

14. How many caffeinated beverages do you have a day? (0 = min, 10+ = max)

15. What is the average time you spend outdoors exposed to direct sunlight on typical day? (Please indicate hours on the left and minutes on the right. For example, if you spend an hour and a half in direct sunlight, please enter 01 on the left and 30 on the right)

*This variable was reverse coded (1 = minimum, 10 = maximum)

**This variable is referred to as sleep inertia throughout the manuscript
